# Diversity without borders: partitioning continuous spaces using probabilistic equivalent numbers

**DOI:** 10.64898/2026.08.10.743903

**Authors:** Pablo Castro Sánchez-Bermejo, Joaquín Hortal, Esben Moland Olsen, Cristina Ronquillo, David Villegas-Ríos, Carlos P. Carmona

## Abstract

Equivalent numbers represent biodiversity as the effective number of equally distinct units, typically species, and can be partitioned across scales. In practice, they summarize each unit of biodiversity by a single value and compare units pairwise, misrepresenting units that are better described as distributions and the relationships between several units that share the same space. We introduce an equivalent-number index for assemblages of units represented as probability density functions (PDFs) over a continuous space, estimated as the integral of the pointwise maximum across abundance-weighted PDFs. Resulting equivalent PDF numbers fulfil elementary properties of classical equivalent numbers, and support additive partitioning across any number of nested scales. We illustrate the framework with case studies across three domains: (1) measuring trait diversity considering intraspecific variability in grasslands, (2) partitioning realized bioclimatic niches among clades of Carnivora, and (3) understanding seasonal changes in the partitioning of fish home ranges in geographic space.

## INTRODUCTION

Many of the entities that ecologists work with are not points but distributions over continuous spaces. Across disciplines as varied as evolutionary ecology, biogeography, movement ecology and functional ecology, representing units as distributions in a continuous space has become standard practice (Carmona *et al*. 2016; Soberón 2007; Worton 1989), supported by growing datasets and tools to estimate these distributions. A species’ climatic niche represents the multidimensional volume of conditions under which it can persist (Hutchinson 1957); an animal’s home range describes the set of locations where it is likely to be found over a given period (Börger *et al*. 2008); and a species’ position in trait space can include the variation among individuals beyond the mean (Palacio *et al*. 2025; Violle *et al*. 2012). Yet the frameworks we use to measure and partition the diversity of assemblages of such units have not kept pace; developed for discrete entities characterized by single values, they translate poorly to this richer representation.

Among the frameworks available to measure diversity, equivalent numbers stand out for their direct interpretation as the effective number of equally distinct units (Hill 1973). An equivalent number is bounded between one and the number of units in an assemblage; it reaches its maximum only when units are equally abundant and fully dissimilar, and tends to one as abundances become uneven or units more similar (Jost 2007). Equivalent numbers have been used to assess taxonomic, functional and phylogenetic diversity (Chao *et al*. 2014), and to build diversity profiles that bridge classical diversity indices through a single parameter tuning the sensitivity to dominance (Leinster & Cobbold 2012). They also fulfil partition properties that allow diversity to be decomposed across hierarchical levels (hereafter “scales”; De Bello et al. 2010; Figure 1a). In practice, however, equivalent numbers have mostly been applied by summarizing each unit through a single value (e.g. a species’ mean trait or niche centroid) and computing pairwise dissimilarities from those summaries. This is a poor approximation when a unit is better represented as a distribution, as is the case for climatic niches (Vásquez-Valderrama *et al*. 2022), home ranges (Worton 1989), and traits with substantial intraspecific variation (Violle *et al*. 2012).

**Figure 1.**
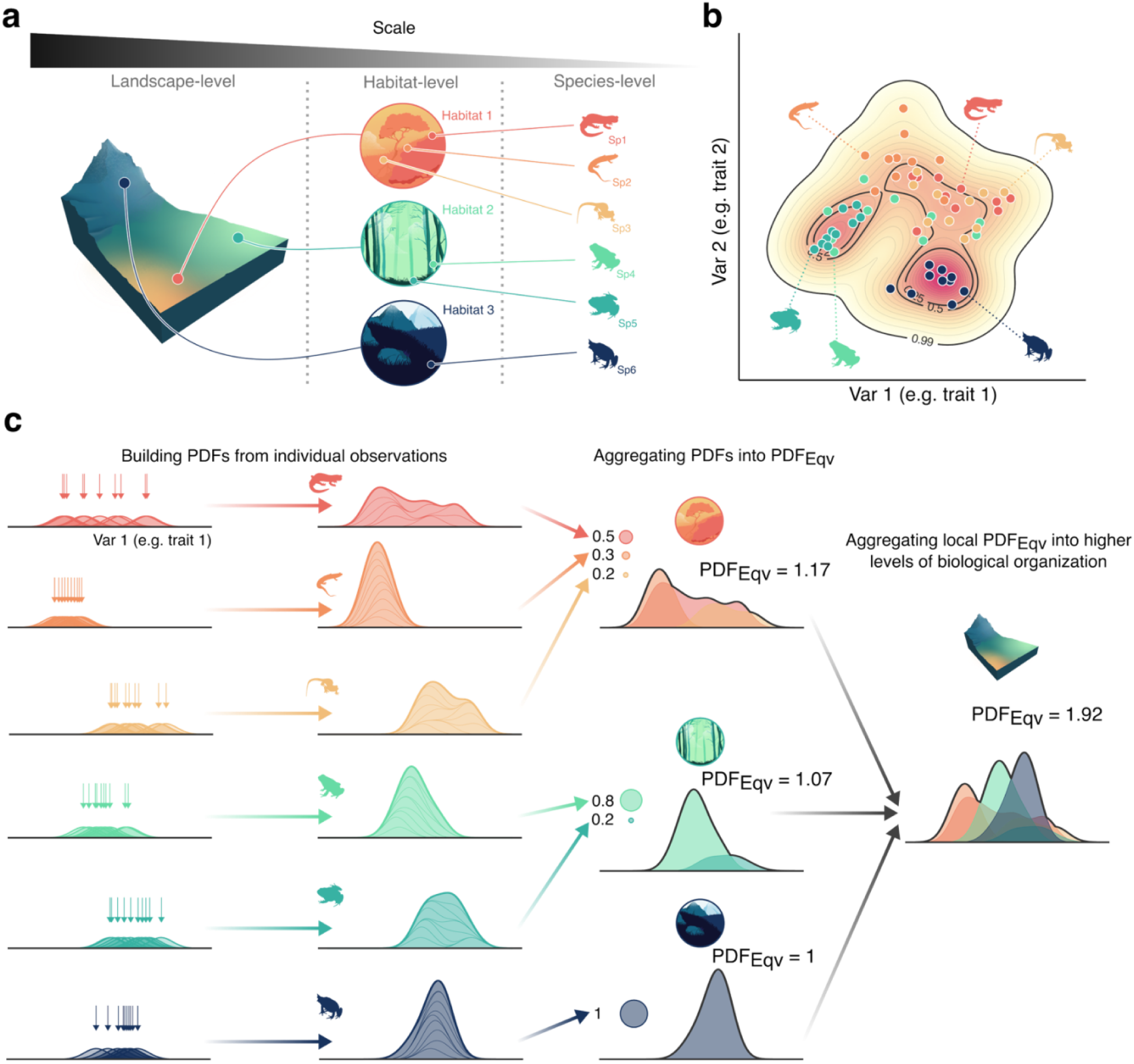
Partitioning the diversity of a continuous space across scales. The organization of diversity across hierarchical scales is exemplified in (a) by showing a nested structure where a landscape is composed of different habitats, each holding different species of amphibians. Observations of each species can be placed in (b) a continuous space representing one facet of diversity (e.g. trait space). Continuing with this example but only using one dimension for clarity, we show (c) how to calculate the equivalent probability density function numbers (PDF_Eqv_). By using probability density functions (PDFs) estimated from individual observations, PDF_Eqv_ can be measured as the integral of the function resulting from the maximum value of the PDFs in an assemblage (black line along the edge of the curves), considering that each PDF integrates to one. In addition, as the relative abundance of the units can be uneven (e.g. census data of species in a plot), the PDFs of each species are rescaled by the abundance of the most abundant species, with the possibility controlling the weight given to rare versus dominant species. PDF_Eqv_ from different assemblages can be combined, allowing to partition diversity additively across any number of hierarchical levels.

An alternative line of work represents ecological units as distributions in a continuous space rather than as point summaries (Carmona *et al*. 2016; Colwell & Rangel 2009; Soberón 2007; Soberón & Nakamura 2009). Probability density functions (PDFs) are a natural choice to represent these units (Figure 1b); they describe how observations are distributed over the space, capturing both extent and shape (Carmona *et al*. 2016; Mason *et al*. 2011). PDFs have been used to estimate animal home ranges (Fleming & Calabrese 2018), to compare a species’ climatic niche across its native and non-native ranges (Vásquez-Valderrama et al. 2022), and to characterize trait diversity in plant and animal assemblages (Mason et al. 2011; Rodriguez-Caro et al. 2023). From PDFs one can also measure the extent (Blonder 2018; Blonder *et al*. 2014) and shape (Carmona *et al*. 2019) of occupied space or quantify pairwise dissimilarity through PDF overlap (Traba *et al*. 2017; Wong & Carmona 2021). These overlap approaches account for the full distribution rather than a single value, yet, being pairwise, they cannot capture how multiple units jointly occupy a space. More fundamentally, PDF-based methods provide no direct equivalent-number measure of total diversity and lack the partition properties that make equivalent numbers useful for decomposing diversity across scales. No framework so far combines the distributional realism of PDFs with the formal structure of equivalent-number indices.

Here, we present an equivalent-number framework for assemblages of continuous-space units, treating each unit as a PDF (Figure 1c). We define the equivalent numbers of PDFs in an assemblage (PDF_Eqv_), show that it shares the elementary properties of equivalent-number indices, and that it supports an additive partition of diversity across any number of nested scales, each contributing a non-negative share to total γ-diversity. We then illustrate the framework with three case studies from distinct domains: trait diversity along a topographic gradient in a Mediterranean grassland; bioclimatic niche partitioning across nested levels of the taxonomic hierarchy of the order Carnivora; and seasonal shifts in the spatial partitioning of Atlantic cod in a Norwegian fjord. We provide the functions needed to implement the method.

## THE EQUIVALENT PROBABILITY DENSITY FUNCTION NUMBERS (PDF_Eqv_)

A probability density function (PDF) describes how observations are distributed over a continuous space; its integral over any region gives the probability mass in that region, and PDFs always integrate to one across the whole space. For each unit in the assemblage, we estimate a PDF from its observations in the continuous space, in the same spirit as the trait probability density (TPD) framework (Carmona *et al*. 2016). Previous TPD-based applications stack unit-level PDFs into a single PDF representing the assemblage, whereas here the PDFs remain separate and enter the diversity calculation through their geometric overlap in the space. The choice of what constitutes a unit is application-specific, and the analytical framework operates the same way regardless of that choice. For example, in the three case studies that follow, the units are populations of plants in a Mediterranean grassland, species of mammals across the Carnivora order, and individual fish in a Norwegian fjord. This choice is preserved at every higher scale of analysis: if units are species, PDF_Eqv_ measures an effective number of species at any aggregation level; if units are individuals, an effective number of individuals; and so on.

We define the equivalent number of PDFs in an assemblage (PDF_Eqv_) as the integral of the pointwise maximum of the PDFs of its units across the space (Figure 1c). We use “assemblage” in a broad sense, to define any set of units grouped together for analyses, be it plant populations within a community, mammal species within a lineage, fish individuals moving along a portion of the sea, or any other set of biological entities sharing a common spatial, ecological, or evolutionary space. Geometrically, PDF_Eqv_ corresponds to the portion of the continuous space effectively occupied by the assemblage: when units occupy disjoint regions their PDFs contribute additively to the integral, and when they overlap the integral counts only the highest contribution at each position. Because in many applications units are not equally abundant in an assemblage, each PDF is rescaled by the relative abundance of its unit, normalized to the abundance of the most abundant unit as:

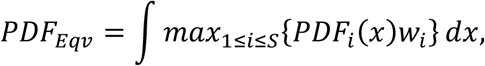

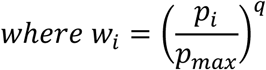

where *S* is the number of units in the assemblage, *PDF_i_*(*x*) is the PDF of unit *i* over the continuous space, *p_i_* is its relative abundance, *p_max_* is the abundance of the most abundant unit in the assemblage, and *q* represents a non-negative number that controls the sensitivity of the index to dominance in a way that is analogous in spirit, though not in form, to the order parameter in classical equivalent-number indices (Jost 2007; Leinster & Cobbold 2012). *w_i_* is the rescaled weight of unit i, which ranges between zero and one, and equals one for the most abundant unit. The parameter *q* sets how steeply the weights of the remaining units decay, so that at *q = 0* all units are treated as equally important regardless of their abundance; as *q* increases, the index depends increasingly on the most abundant units, and rare ones contribute less. This makes it possible to study how assemblage diversity varies across the dominance spectrum by describing how PDF_Eqv_ changes with q (Leinster & Cobbold 2012). Although PDF_Eqv_ is built from a different mathematical structure than classical equivalent numbers, which are derived from entropy-based estimations (Hill 1973), we show that it shares their key elementary and partition properties for any value of q (see Box 1 and Figure 2).

**Figure 2.**
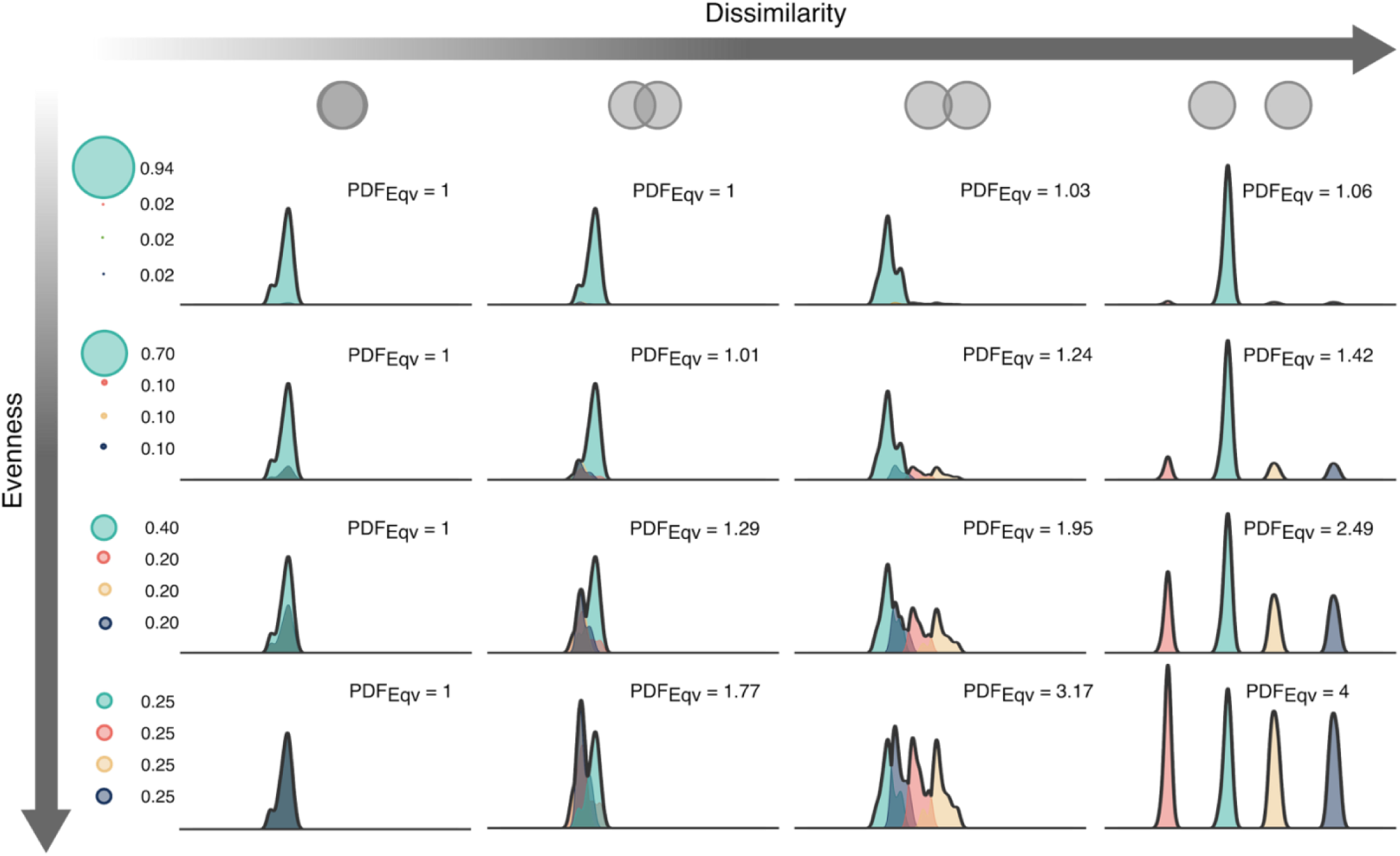
Changes in probability density function equivalent numbers (PDF_Eqv_) with dissimilarity and evenness, in a simulated assemblage composed of four units. Each distribution is the probability density function (PDF) of a unit, with one colour per unit. Dissimilarity increases from left to right and evenness increases from top to bottom. PDFs are rescaled relative to the most abundant unit. The black line over the PDFs is the pointwise maximum of the rescaled PDFs, and its integral is PDF_Eqv_. In the equal abundance scenario (bottom row), PDF_Eqv_ depends solely on overlap: it equals the number of units when they are fully dissimilar (bottom right corner), and decreases toward one as overlap increases (moving left). Conversely, in the highly uneven scenario the weight of the non-dominant units, computed relative to the most abundant one, are small; PDF_Eqv_ therefore tends to one with little effect of overlap among units (top row).

### BOX 1. Elementary properties of PDF_Eqv_s

Whether an index captures the desirable features of diversity is evaluated by examining a set of properties (Hubálek 2000). Equivalent numbers must fulfil several elementary properties, related to the range of values and its sensitivity to the number of units, together with partition properties for decomposing diversity across scales (see Appendix S1 for partition properties). We evaluate five elementary properties that Pavoine (2026) used to assess equivalent numbers: (1) non-negativity, (2) minimum value under total unevenness or full similarity, (3) maximum value under total evenness and full dissimilarity, (4) sensitivity to the number of units when units are evenly represented and fully dissimilar and (5) weak monotonicity.

First, PDF_Eqv_ integrates the pointwise maximum of weighted PDFs, all of which are non-negative; the index is therefore non-negative as well. The second property concerns the lower bound: the minimum value should be reached when units are fully similar. As illustrated in Figure 2 (left column), if *S* units are fully similar, so that ∫ *min* {*PDF_i_* (*x*), *PDF_j_*(*x*)}*dx* = 1, then all PDFs coincide; the maximum function is this common PDF multiplied by the largest weight, which is one, so its integral is one:

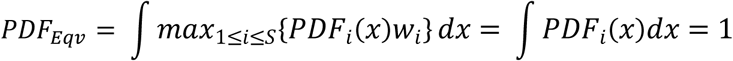

The minimum is also reached when the weights are totally uneven so that a single unit carries all the weight (as in an assemblage of one unit; Figure 2 top row). In this case, the maximum function is simply the PDF of the single unit:

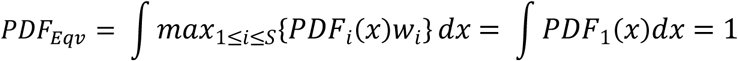

More generally, when q > 0 the weights of the less abundant units shrink and one unit becomes more dominant, so the index declines toward one (Figure 2, top row).

Conversely, the upper bound is the number of units present in the assemblage (Hubálek 2000). The third property requires this maximum to be reached when abundances are even and units are fully dissimilar. If all units are equally abundant every weight equals one (*w_i_* = 1 for any value of q, since *p_i_* = *p_max_* for all i), and the index becomes the integral of the maximum of the unweighted PDFs:

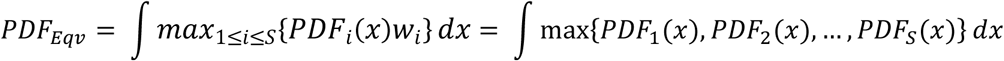

and considering that ∫ min{*PDF_i_*(*x*), *PDF_j_*(*x*)}*dx* = 0 for any *i* and *j*, then:

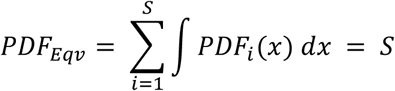

Because the integral of each PDF equals one, the value of the index equals the number of units (S) in this case (Figure 3 bottom right panel). Further, given that under the conditions of even abundance and full dissimilarity the index equals the number of units, for two assemblages with S_1_ and S_2_ units, where S_1_ > S_2_, then PDF_Eqv1_ > PDF_Eqv2_, which meets the fourth property about sensitivity to the number of units.

Last, monotonicity is the property of a diversity index never decreasing when a new unit is added to an assemblage (Hubálek 2000). Strict monotonicity can fail for indices that account for abundances; the relevant requirement is therefore weak monotonicity, which means the index does not decrease when a unit no more abundant than those already present is added (Pavoine 2026; Weikard *et al*. 2006).

At q = 0, PDF_Eqv_ is presence-based: abundances drop out, so adding a unit can only enlarge the maximum function, and strict monotonicity holds trivially. The property is only at stake for q > 0 where the PDFs are rescaled relative to the most abundant unit.

The case q > 0 needs more care, because PDFs are rescaled relative to the most abundant unit. As long as the added unit is not the most abundant, the reference is unchanged and the index cannot decrease. Labelling the most abundant unit as 1, for an assemblage of S units with abundances *p* = (*p*_1_, …, *p_S_*), and *p*_1_ = *max*(*p*):

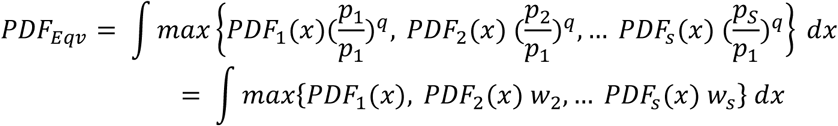

Adding a unit with PDF_new_ and abundance *p_new_* < *p*_1_ leaves *p*_1_as the reference, so the existing weights are unchanged; the new term 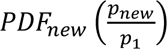 is an additional function inside the maximum, which can only raise or leave the index unchanged.

The situation differs when the added unit is more abundant than every unit already present (p_new_ > p1). It then becomes the new reference and the weight of every existing unit is recomputed relative to p_new_:

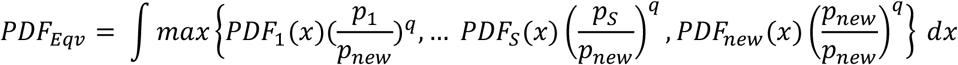

Because every weight is not smaller, the existing units contribute less to the maximum function, and depending on how the rescaled PDFs overlap, the index can fall below its previous value. Therefore, PDF_Eqv_ can decrease only when the added unit is more abundant than the current maximum, fulfilling the property of weak monotonicity.

### Partitioning of diversity based on PDF_Eqv_s

Equivalent numbers have been particularly influential for partitioning diversity across hierarchical scales, separating the diversity within assemblages from the contributions of differences between them (de Bello et al. 2010; Jost 2007). For PDF_Eqv_ we use an additive partition, which is natural in our framework because the maximum function aggregates as a geometric union, so that the diversity contributed by each scale corresponds to the additional portion of the space unlocked at that scale. To aggregate across scales, we perform a maximum-of-maximum operation by taking the pointwise maximum of the within-assemblage maximum functions at each higher level, without re-scaling abundances at the new scale (Figure 1c). The pointwise maximum is at least as large as each individual maximum function, so γ ≥ ᾱ at every level of aggregation and the additive between-assemblage component β = γ − ᾱ is non-negative. Leinster & Cobbold (2012) suggested that equivalent numbers should fulfil three partitioning properties: (1) sensitivity to the number of units when species are evenly represented and fully dissimilar, (2) modularity and (3) replication. These properties are proved in Appendix S1. This partition extends to any number of nested scales (e.g., populations within species within clades, or quadrats within plots within sites; Carmona et al. 2016), with each scale contributing a non-negative share to the total γ-diversity (Figure 1c). For instance, PDFs representing the climatic niche of several populations of the same species can be aggregated into a single PDF_Eqv_ that summarizes the climatic niche of the species (expressed in equivalent populations), and this procedure could be repeated across higher-order scales.

## CASE STUDIES

To illustrate this framework, we present three study cases from different ecological and evolutionary domains. In the first, we measure trait diversity in a Mediterranean grassland from PDFs capturing intraspecific variation in two functional traits and examining how trait dimensionality affects the measured partition. In the second, we evaluate the bioclimatic niche partition of the order Carnivora across nested levels of the taxonomic hierarchy to assess phylogenetic niche conservatism. In the third, we quantify how individuals of Atlantic cod partition the geographic space of a Norwegian fjord across seasons. Together, these cases span functional ecology, biogeography, and movement ecology.

### Case study 1: Trait partitioning and dimensionality in a Mediterranean grassland

We aim to understand how dimensionality influences the partition of the trait space in a Mediterranean grassland. Equivalent numbers offer a natural measure of diversity interpretable in terms of trait partitioning; through diversity profiles, they also can reveal how strongly dominance shapes that diversity (Leinster & Cobbold 2012). In practice, most studies applying equivalent numbers consider pairwise comparisons of mean species trait values (De Bello *et al*. 2010; Chao *et al*. 2014; Jost 2007) using Gower’s dissimilarity, the average of scaled dissimilarities across individual traits (Gower 1971). This approach misses two fundamental features. First, by reducing species to a mean, it neglects within-species variability, an important component of trait diversity (Castro Sánchez-Bermejo *et al*. 2025; Wong & Carmona 2021). Second, because Gower’s dissimilarity averages across dimensions, adding a trait on which species are similar lowers their mean dissimilarity, even though in the trait space itself two species can only grow more separated, never less, as more traits are considered (Maire *et al*. 2015). Using PDF overlap as the dissimilarity metric addresses both problems, since it uses the full distribution in the trait space (Bello *et al*. 2013); yet it remains pairwise, and a matrix of pairwise overlaps cannot capture how all units jointly occupy that space. PDF_Eqv_s removes this limitation by operating on the geometric union of all PDFs through the pointwise maximum, capturing the joint occupation of the trait space directly. Here, we compare diversity profiles from PDF_Eqv_ and the Hill-Chao framework with Gower’s dissimilarity, across trait spaces built from one and two traits, to test whether accounting for the joint occupation of trait space changes how dimensionality affects measured diversity.

We used individual-level data for height and specific leaf area (SLA) across 40 plots in a Mediterranean grassland (see Method S1 for details). With the *TPDs* function in the *TPD* package (Carmona *et al*. 2019) we estimated PDFs for each trait independently and for both traits together, per species and plot. We calculated PDF_Eqv_ for each plot accounting for each species’ relative abundance. For the comparison, we averaged trait values and computed standardized dissimilarities for SLA and height, combining both traits with Gower’s distance (FD package; Laliberté et al. 2014), and used them to calculate equivalent numbers under the Hill-Chao framework (Chao *et al*. 2014). A common value for ‘τ’, the threshold of functional distinctiveness which determines if differences between species are considered meaningful (Chao *et al*. 2014), was calculated as the average abundance-weighted mean pairwise functional distance calculated for SLA, height and the Gower’s dissimilarity to allow comparisons. In both frameworks, equivalent numbers were calculated from q = 0 (rare and abundant species weighted equally) to q = 2 (emphasizing the weight of abundant species), in steps of 0.1. For each q and framework, we compared equivalent numbers across the trait space used (height, SLA, or both) through linear mixed-effects models, with the trait space as a fixed factor and plot position along the topographic gradient as a random effect (Kuznetsova *et al*. 2020). Significance was assessed by likelihood-ratio tests, with Tukey post-hoc tests between spaces.

We found that the number of equivalent species (PDF_Eqv_) increased with dimensionality, being higher in the two-dimensional trait space than in either single-trait space (Figure 3; Table S1). This increase was expected, since adding a trait axis can only preserve or increase the separation between species in the joint space. (Barry *et al*. 2019). This dimensionality effect remained significant even though the range of PDF_Eqv_ values narrowed with q (Figure S1), which indicates that most communities are dominated by a few species. By contrast, the Hill-Chao framework with Gower’s distance showed the opposite: no significant differences between trait spaces at any value of q (Table S2). Because Gower averages dissimilarity across traits, a genuine separation on one axis is diluted by similarity on another, and the dimensionality signal that PDF_Eqv_ recovers is cancelled. The two frameworks thus encode different notions of dissimilarity; when trait axes represent genuine independent dimensions, preserving their joint separation, as PDF_Eqv_ does, reflects how species partition the trait space more faithfully than averaging across axes. Two plants alike in one trait but well separated by another (Barry et al. 2019) are counted as distinct by PDF_Eqv_ and largely merged by Gower.

**Figure 3.**
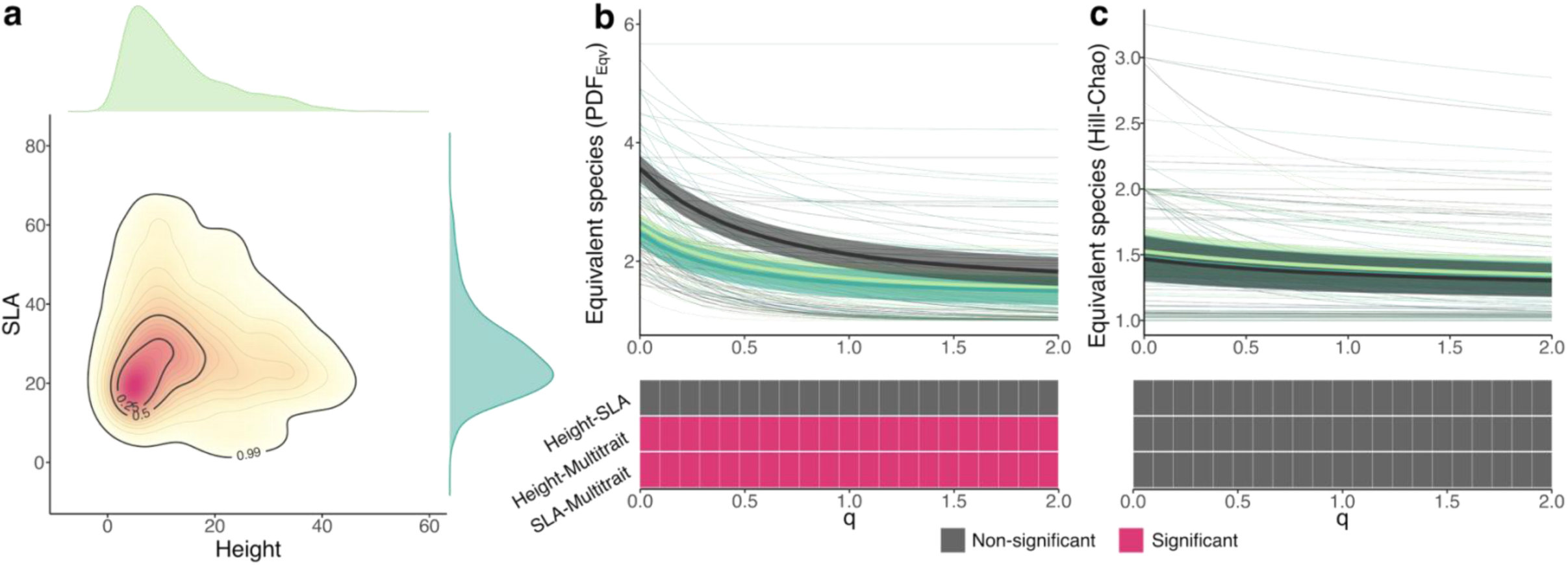
Effects of the continuous space used on the number of equivalent species calculated as equivalent probability density function number (PDF_Eqv_) and using a Hill-Chao framework. Equivalent numbers were calculated using plant height (light green), specific leaf area (SLA; dark green) and the two traits together (black) on 40 plots sampled along a topographic gradient in a Mediterranean grassland under different values of a parameter ‘q’ which weights importance of the abundance of the species in the calculation of PDF_Eqv_. The density plots in **(a)** show the aggregation of PDFs from species and communities as in Carmona et al. (2016). The colour gradient visualizes different probability densities of the two-dimensional continuous space, and portions of the space with highest densities of observations correspond to red colours. Diversity profiles ranging between q = 0 and q = 2 were calculated for each plot using **(b)** a PDF_Eqv_ and a **(c)** Hill-Chao framework (thinner lines). Linear mixed-effects models were fitted for each value of q (at a resolution of 0.1). Thicker lines and shaded areas represent model estimates and confidence intervals. Using likelihood ratio test we found that the equivalent species differed significantly depending on the continuous space used **(b)** across all values of q, for which Tukey tests revealed that the number of equivalent species was higher (p < 0.001 in every case; see Table S2) in the case of the two-dimensional continuous space compared to the continuous spaces built using one variables (SLA or height). In contrast, we did not find any difference between the number of equivalent species using SLA-based dissimilarities, Height-based-dissimilarities or Gower’s based dissimilarities.

### Case study 2: Patterns of bioclimatic niche conservatism in Carnivora

We aim to understand how the bioclimatic niche is partitioned across lineages. Evolutionary processes tend to leave closely related species with more similar niches than distant relatives (phylogenetic niche conservatism; Hadly et al. 2009; Pyron et al. 2015), a pattern usually studied through niche descriptors (e.g. mean and variance; Olalla-Tárraga et al. 2011) or pairwise overlap between species (Peixoto *et al*. 2017). Here we estimate climatic niche conservatism with PDF_Eqv_, computing each species’ PDF in a climatic space and aggregating them along the clades of Carnivora. For each clade, PDF_Eqv_ ranges from 1 (no partitioning), to the number of species in the clade (total partitioning). Carnivora is a good study model because it shows well-known adaptations to different habitats, from the marine environment of pinnipeds (Berta *et al*. 2018) to the divergent climates occupied by closely related species, such as the genus *Vulpes* (Kumar *et al*. 2015). Such diversity of adaptations should translate into contrasting patterns of niche partitioning across groups. We assess which clades show niche conservatism (fewer equivalent species than expected by chance) and which show divergence (equivalent species close to the number of species in the clade), testing whether divergence is concentrated in clades that adapted to novel climates.

We retrieved all Carnivora occurrence records available at Global Biodviersity Information Facility (GBIF.org 2026; 7,940,261 records) using the *rgbif* package (Chamberlain *et al*. 2025), then validated and filtered them following Ronquillo et al. (2024; see Method S2 for details). Occurrences were combined with six bioclimatic variables and used to build a PCA where we retained two principal components (climatic space; Figure 4a; Method S2). We used the *TPDs* function (Carmona *et al*. 2019) to characterize the climatic niche of every species as a PDF. We assumed evolutionary relationships between taxonomic groups following a recent phylogeny of Carnivora inferred from mitochondrial genomes (Hassanin *et al*. 2021). PDF_Eqv_ was calculated for each genus, subfamily, family, infraorder (Arctoidea, Canoidea, Viverroidea and Feloidea) and suborder (Caniformia and Feliformia). Afterwards, observed PDF_Eqv_ values were compared against 500 null expectations in which species identities were randomized across taxonomic groups, keeping the number of species within each group fixed. These null models provide a baseline of no climatic niche conservatism; deviation from this expectation was measured with standardized effect sizes (SES; Gotelli & McCabe 2002), and observed PDF_Eqv_ were considered significant when they fell outside the 95% confidence interval of the simulated values.

**Figure 4.**
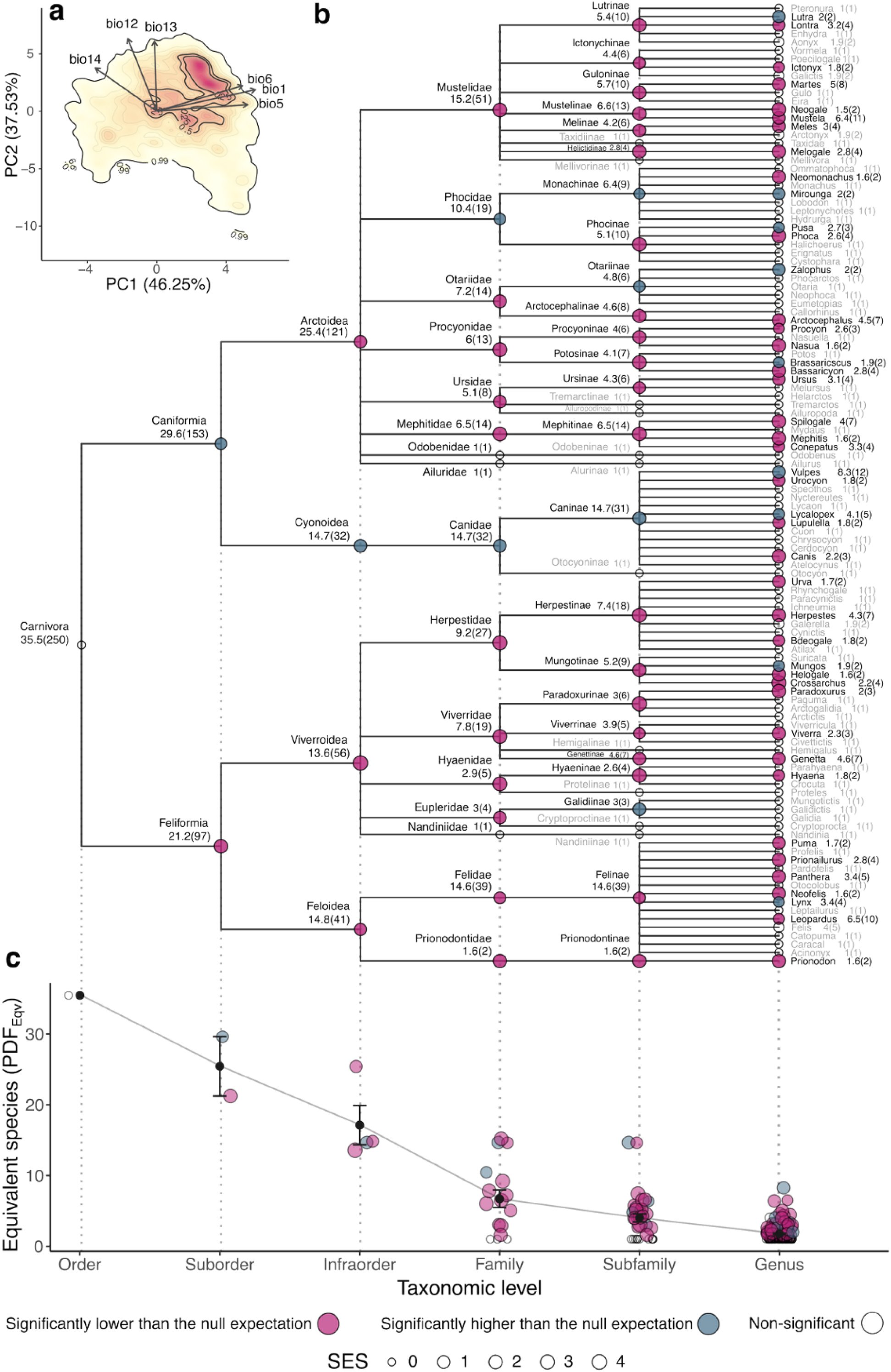
Equivalent number of species in the order Carnivora, calculated as equivalent probability density function numbers (PDF_Eqv_). (a) Cladogram showing the equivalent number of species (with total number of species in parenthesis) for each taxonomic groups across levels. Points indicate the deviation from the null expectation for each taxonomic level: size = absolute standardized effect size (SES); colour = positive (blue) or negative (pink) deviation. Text colour indicates significance: grey = no significant deviation from the null expectation, black = significant deviation. (b) Scatter plot of differences in equivalent numbers across taxonomic levels; black points and error bars indicate the mean and standard error per group.

Considering all extant Carnivora together, the number of equivalent species in the bioclimatic space was 35.3 (Figure 4b), decreasing towards lower taxonomic levels until reaching 1 at the species level (Figure 4c). Far from being constant, this decrease is steeper at higher levels (e.g. from Order to Suborder) than at lower ones (e.g. from Family to Genus), suggesting that closely related species tend to occupy similar areas of the climatic space. The null models support this: among clades deviating significantly from the null expectation, PDF_Eqv_ was mostly lower than expected (36 of 45 genera, 20 of 24 subfamilies, 11 of 13 families, 3 of 4 infraorders and 1 of 2 suborders). Species within a taxonomic group therefore tend to partition their bioclimatic niche less than expected by chance, retaining climatic adaptations from a common ancestor so that close relatives remain ecologically more similar than distant ones (e.g. Olalla-Tárraga et al. 2011; Pyron et al. 2015; Wiens & Graham 2005). In other words, niche conservatism is the norm among Carnivora (e.g. genus *Crossarchus*; see Figure S2; Cooper et al. 2011; Liu et al. 2020)).

Some taxa depart from this pattern, showing greater partitioning than expected by chance: nine genera (*Lutra*, *Mirounga*, *Pusa*, *Zalophus*, *Brassariscus*, *Vulpes*, *Lycalopex*, *Mungos*, *Lynx*), four subfamilies (Monachinae, Otariinae, Caninae, Galiidinae), two families (Phocidae, Canidae), one infraorder (Cyonoidea), and one suborder (Caniformia). The genus *Vulpes* is a good illustration, since it covers most of the bioclimatic spectrum, and except for some generalist species such as *V. vulpes*, its species are highly climatically specialized (e.g. *V. rueppellii* and *V. zerda* in deserts, *V. lagopus* in the Arctic; Figure S3), indicating strong niche divergence. Such divergence is expected under ecophysical allopatry, where physical and environmental heterogeneity contribute to speciation (Pyron *et al*. 2015), which may explain why species in divergent groups occupy distinct geographical distributions (e.g. Phocidae; Fyler et al. 2005). We found evidence for patterns of both niche conservatism and divergence at the higher taxonomic levels, allowing to trace the signal of past climate adaptation back to lineages that split in the Eocene and Miocene (Feliformia and Caniformia, respectively; Hassanin et al. 2021): Feliformia shows low niche partitioning, evidencing strong conservatism, whereas Caniformia shows high partitioning across most of its evolutionary history, driven by adaptation to novel climates in Phocidae and Canidae (Figure 4b).

### Case study 3: Movement of Atlantic cod in a Norwegian fiord

We used an acoustic telemetry dataset of 25 individuals of Atlantic cod (*Gadus morhua*) from the Tvedestrand fjord, on the Norwegian Skagerrak coast (Villegas-Ríos et al. 2020; Figure 5a), to understand how fish partition geographic space throughout the year. Atlantic cod shows strong seasonality in movement, driven mainly by their mating system (Zemeckis *et al*. 2017), which resembles a lek: males defend spawning territories that female visit during the spawning season. We study the partitioning of geographic space through PDF_Eqv,_ treating each individual’s home range as a PDF (Worton 1989), and test the hypothesis that during the spawning season cod partition the fjord more markedly than during the rest of the year.

**Figure 5.**
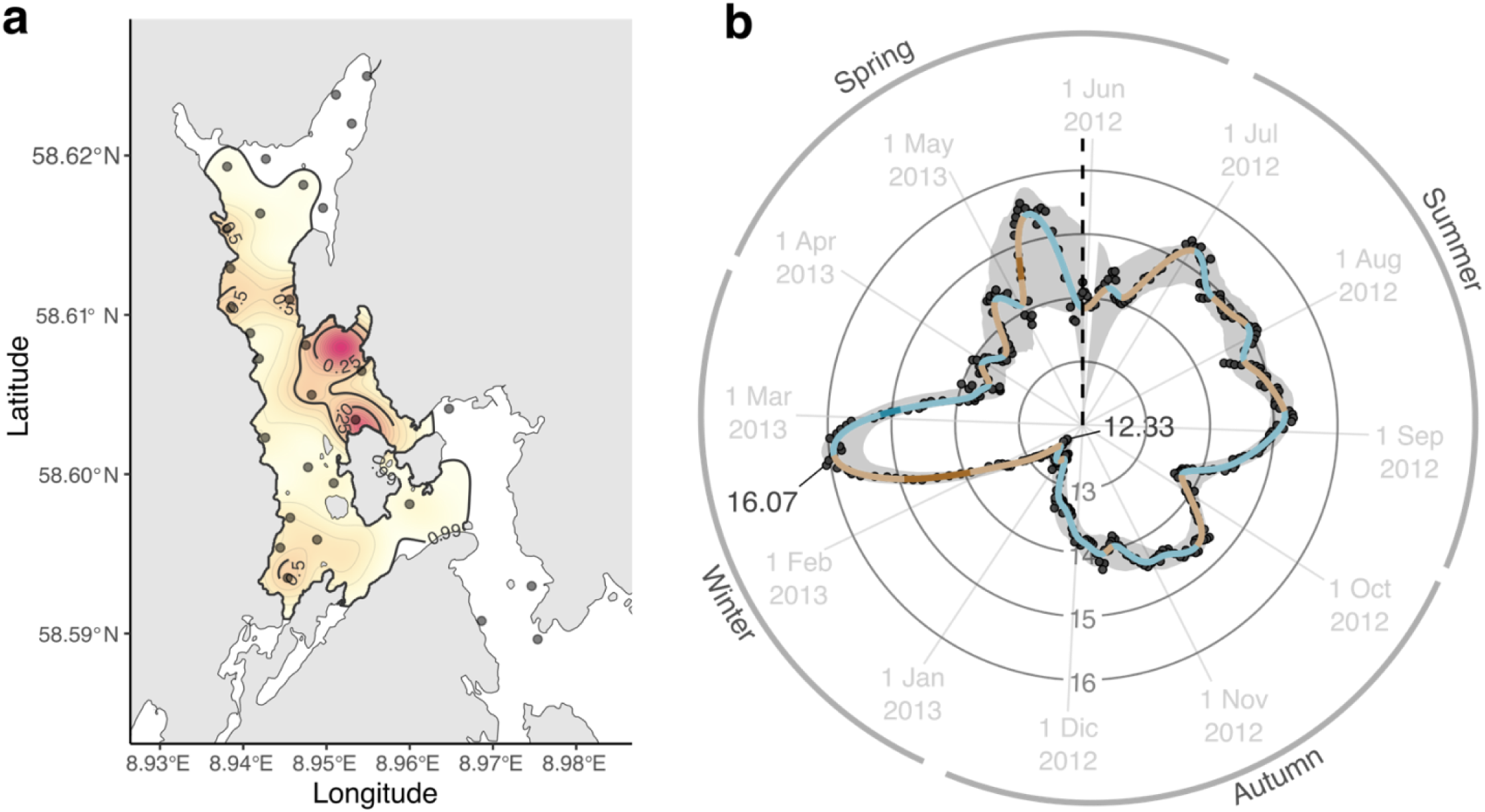
Seasonal changes in the partition of the geographical space in a Norwegian fjord. The partitioning of the (a) occupied geographical space Tvedestrand fjord, on the Norwegian Skagerrak coast, by 25 individuals of Atlantic cod (*Gadus morhua*) along one full year using a sliding-window approach which considered the occupied space by individual fish during a two-week period. The density plot in (a) shows the aggregation of the probability density functions (PDFs) for all fishes in the whole year (see Figure S5), with colour gradient showing the estimates of the function, and red colours corresponding to portions of the geographical space with highest densities of observations. Points show the location of the receivers in Tvedestrand fjord. The circular plot displayed in (b) show the fluctuations during a whole year in the equivalent number of home ranges (calculated as equivalent probability density function number (PDF_Eqv_)), with increases (brown) and decreases (blue) along the year. Significant trends, as revealed by a derivative analysis, were found in February, March and May (solid colours). The dashed line indicates the beginning and end of the time series, while the lines to the points indicate the maximum and minimum values.

The dataset encompassed 25 fish monitored between May 24^th^ 2012 and June 5^th^ 2013 (see Method S3 for details). To track changes in the use of the geographic space throughout the year, we used a sliding-window analysis with two-week windows, wide enough to ensure sufficient data to estimate home ranges for each individual. Within each time window, all centres of activity (COAs) per fish were used to estimate its home ranges as a PDF in geographic space (longitude and latitude) via the *density.ppp* function of the *spatstat.explore* package, applying an edge correction to avoid boundary problems, and a bandwidth calculated per individual and window with the *kde* function of the *ks* package (Duong 2024; Mcswiggan 2026); PDF_Eqv_ then gives the equivalent number of home ranges per window (ranging between 1 and 25). The window was advanced one day at a time, adding the next COAs and dropping the previous ones. The procedure was repeated 365 times to cover a full year. Each window contained on average 593.46 COAs per individual (minimum = 89; maximum = 672). Trends across the year were modelled with generalized additive models (GAMs) using a cyclic spline to account for seasonality (Perperoglou *et al*. 2019) and a basis dimension (k) of 50 (Pedersen et al. 2019; see Figure S4). To handle the temporal autocorrelation introduced by overlapping windows, we used a moving block bootstrap (Ju 2015), resampling with replacement consecutive 7-day blocks to preserve the temporal dependence structure. We fitted one GAM per bootstrap iteration (500 in total), and estimated uncertainty in the seasonal pattern from the bootstrap distribution of fitted curves. Trends in the average estimate were assessed by calculating the derivatives and their standard errors at each point, taking increases or decrease as significant when the confidence interval excluded zero (Simpson 2018).

We observed drastic changes in the way cod individuals partition the fjord, which coincide with seasonal patterns of the behaviour of the species. The equivalent number of home ranges varied throughout the year, between 12.33 and 16.07 (Figure 5b), indicating that the degree to which cod partition the fjord differs between periods. Summer and autumn 2012 showed no significant trends despite slight fluctuations. In contrast, partitioning rose steeply during winter, going from the lowest to the highest yearly values within a month (12.33 on January 9^th^ 2013; 16.07 on February 12^th^ 2013). This coincides with the onset of the spawning season around February (Cianelli *et al*. 2010) and would reflect males arriving first at spawning sites, where they defend territories (Dean *et al*. 2014; Skjæraasen *et al*. 2026). Partitioning then decreased significantly, remaining around 13 to 14 from mid-March until May 2013, close to summer and autumn values. This decrease may reflect females moving among male territories (Olsen *et al*. 2023), consistent with a bet-hedging strategy of maturing and spawning batches of eggs at various locations over several weeks (Hočevar *et al*. 2021). A second steep and significant increase followed in May 2013, when spawning is most likely over and behaviour shifts from spawning to feeding (Zemeckis *et al*. 2017). At this time, cod disperse to feeding habitats across the fjord (Freitas *et al*. 2021), so the rise in partitioning probably indicates divergence in feeding habitat use, in line with stable isotope evidence of diverse feeding behaviours and diets within this population (Monk *et al*. 2023). By the end of May 2013, values of partition were similar to those at the start of the time series in May 2012.

## CONCLUDING REMARKS

Here we show that the same formal structure describes how a community partitions its trait space, how a clade partitions its bioclimatic niche, and how a population partitions its habitat through time. The disciplines represented in these case studies already represent their units as distributions, but they lacked a common way to express the diversity those distributions imply; PDF_Eqv_ provides such a common metric, and should make comparative work across disciplines more straightforward.

Our three case studies illustrate the use of the framework, but they are far from taking its whole profit. Because PDF_Eqv_s can be calculated at any level of biological organization, the applications within ecological disciplines go beyond the case studies presented here: partitioning a species’ trait space by considering variability within individuals (Herrera 2024), comparing the climatic niche of populations of the same species across biogeographical realms (Broennimann *et al*. 2007), or characterizing how different populations or species share the same geographical space (Willems & Hill 2009). More broadly, PDF_Eqv_ applies wherever diversity is measured over continuous variables. Spectroscopy is one example: reflectance metrics measured across wavelengths (Cavender-Bares *et al*. 2022) could benefit from this framework, particularly when spectral data, often composed of hundreds or thousands of measurements are summarized into a continuous spectral space (Beccari *et al*. 2024; Li *et al*. 2023). Similarly, continuous spaces can be used to summarize ecoacoustics recordings (i.e. acoustic trait space; Luypaert et al. 2022) and estimate PDFs (Bacquelé *et al*. 2026). This allows, for instance, using repeated ecoacoustic data from the same location to monitor changes in biodiversity as the equivalent number of recordings.

PDF_Eqv_ brings together two lines of work in ecology. Classical equivalent numbers have been central to understanding taxonomic, functional, and phylogenetic diversity (Chao *et al*. 2014), and have since been extended to other facets of ecosystems, such as ecosystem functions (Chao *et al*. 2024). The use of dissimilarities lends these approaches versatility, allowing them combine categorical and continuous data. They are, however, based on a single value per unit, whereas modern ecology requires tools that allow synthesizing growing volumes of data to describe diversity (Hampton *et al*. 2013). PDFs, by contrast, summarize the occupation of a continuous space realistically even for many observations (Stine & Heyse 2001), which suits them to animal tracking data (our third case study) or to the large number of species occurrence records available from public repositories (our second case study), among other data-intensive applications. Yet PDFs have not offered a direct route to diversity. The TPD framework (Carmona *et al*. 2016) synthesizes them through indices focused on the geometry of the distribution; these indices characterize breadth or shape well, but they lack the formal guarantees expected of diversity indices, such as monotonicity. Approaches based on pairwise overlaps (Mason *et al*. 2011) reduce the occupation of the space to a dissimilarity matrix, which cannot capture how units jointly occupy it. Against this backdrop, we show that expressing PDFs as equivalent numbers through the integral of the maximum function fills this gap: PDF_Eqv_ fulfils the defining properties of equivalent numbers while yielding a diversity metric easily interpretable directly as the partitioning of a continuous space.

The framework has limitations. Some are inherent to PDFs: their estimation requires a bandwidth, the width of the “bell” placed around each observation, which determines the smoothness and therefore the shape of the resulting curve. Different bandwidths can produce different results so bandwidth selection should be approached carefully (Tordoni *et al*. 2024). The computational cost of calculating PDFs increases exponentially with dimensionality (Mammola & Cardoso 2020), which may limit implementation on standard equipment in high-dimensional spaces. Other limitations are inherent to incorporating abundances. Rescaling by *pmax* allows abundances to be incorporated into the PDF_Eqv_ framework, and it is the only reference that preserves the bounds between one and the number of units. As in any abundance-weighted index, this comes at the cost of strict monotonicity: adding a new unit that becomes the new dominant one rescales every weight and can potentially lower the index, so PDF_Eqv_ fulfils weak rather than strict monotonicity (see Box 1; Weikard et al. 2006). The same happens with classical equivalent numbers, which may decrease when a strongly dominant species is added. Finally, PDF_Eqv_ is restricted to continuous spaces, so classical approaches remain valuable where dissimilarities are intrinsically pairwise, as in phylogenetic diversity (Ives & Helmus 2010), or where units are described by categorical variables (e.g., plant phenotype as “grass,” “forb,” or “tree”).

To facilitate the application of the PDF_Eqv_ framework, we developed a set of R functions implementing its main analytical steps, publicly available on Zenodo (https://doi.org/10.5281/zenodo.21870972). The current version includes *PDFeqv*, for calculating PDF_Eqv_ in assemblages, and *diversity.profile*, for calculating and plotting PDF_Eqv_across values of *q*. Both functions take as input observations in a continuous space together with information about the unit identity and an abundance matrix (or a presence/absence matrix when abundance data are not used). Alternatively, an object generated using the TPD package (Carmona *et al*. 2019) can be provided instead of providing individual observations. Two further functions work with objects created by the *PDFeqv* function: *plot.PDFeqv* allows visualization of PDF_Eqv_s, in spaces of up to two dimensions, and diversity.*partition* aggregates PDF_Eqv_s across higher hierarchical scales and partitions it accordingly. Example R code is available alongside this manuscript (https://doi.org/10.5281/zenodo.21870972).

As denser, more multidimensional ecological data become more accessible, methods that operate directly on the distributions of units—rather than on point summaries—will become increasingly central. Beyond the specific cases presented here, we expect the PDF_Eqv_ framework to be most useful in disciplines where ecological units are already represented as continuous distributions but have so far lacked an equivalent-number-style index capable of capturing the diversity these distributions imply.

## DATA AVAILABILITY STATEMENT

All data and R code to reproduce analyses, as well as the R functions developed for implementing the PDF_Eqv_ framework are archived on Zenodo (https://doi.org/10.5281/zenodo.21870972).

## AUTHOR CONTRIBUTIONS

Conceptualization: CPC, with support from PCS-B.

Methodology: PCS-B, with support from CPC.

Software: PCS-B, with support from CPC.

Data curation: CR performed the Carnivora data curation.

Formal analysis: PCS-B, with support from JH, CR, DV-R and CPC.

Visualization: PCS-B.

Supervision: CPC.

Writing—original draft: PCS-B, with support from CPC.

Writing—review & editing: PCS-B, JH, EMO, CR, DV-R, JH.

## ACKNOWLEDGEMENTS

This research was supported by the European Research Council (ERC) under the European Union’s Horizon Europe research and innovation programme (grant agreement No 101126117, PLECTRUM), the Galician Innovation Agency through the Oportunius Program (OTR16264), and the MICIU through the ATRAE Program 2024 (ATR2024-154934). DV-R was supported by a Ramón y Cajal Contract (RYC2021-032594-I), funded by MICIU/AEI and the European Union Next Generation EU/PRTR. CR was funded by Project NICED, grant PID2022-140985NB-C21 by MICIU/AEI/ 10.13039/501100011033 / FEDER, EU. The authors used Claude Sonnet 5 (Anthropic) to assist with improving the readability, internal consistency, and stylistic clarity of the R functions used in this study.

