## Supporting information for "Diversity without borders: partitioning continuous spaces using probabilistic equivalent numbers"

**Appendix S1.** Partition properties.

**Method S1.** Plant functional trait database.

**Method S2.** Carnivora data curation and construction of climatic space.

**Method S3.** Atlantic cod data description and curation.

**Figure S1.** Diversity profiles calculated for 40 plots along a topographic gradient in a Mediterranean grassland using three different continuous spaces.

**Figure S2.** Bioclimatic niche of species in the genus *Crossarchus*.

**Figure S3.** Bioclimatic niche of species in the genus *Vulpes*.

**Figure S4.** Distribution of the (a) effective degrees of freedom and (b) K values for each of the bootstraps in the generalized additive model (GAM).

**Figure S5.** Occupation of the geographic space by each individual in the period of one year.

**Table S1.** Summary of post-hoc Tukey tests comparing the equivalent probability density function number ($PDF_{Eqv}$) calculated using three different trait spaces, two of them based on unique traits (height and specific leaf area (SLA)) and one being a multidimensional trait space formed by these two traits.

**Table S2.** Summary of post-hoc Tukey tests comparing the equivalent numbers (calculated using the Hill-Chao framework) with three different dissimilarity matrices, two of them based on unique traits (height and specific leaf area (SLA)) and one being the Gower's distance using these two traits.

### Appendix S1. Partition properties

According to Leinster & Cobbold (2012), equivalent numbers should fulfil three properties to guarantee partitioning: (1) sensitivity to the number of units when species are evenly represented and fully dissimilar, (2) modularity and (3) replication. We have already seen how the first property is fulfilled in the case of of $PDF_{Eqv}$ (see Box 1), so we now focus on the properties of modularity and replication. Modularity refers to the additive property of equivalent numbers, meaning that in the case of two assemblages with units from different assemblages being completely different, the global diversity is entirely determined by the diversities of the assemblages. In the case of $PDF_{Eqv}$, the global diversity is determined by the maximum value of the $PDF_{Eqv}$s functions assessed at the immediately lower level of biological organization. In the case of several assemblages where:

$\int \min\left\{\int max_{1\leq i\leq S_j}\{PDF_i(x)w_i\}\, dx, \int max_{1\leq i\leq S_k}\{PDF_i(x)w_i\}\, dx\right\}\, dx = 0$ for all $j \neq \mathrm{k}$, then:

$$PDF_{Eq_\Upsilon} = \int max\left\{max_{1\leq i\leq S_c}\{PDF_i\ (x)\ w_i\}\right\} dx = \sum_{j=1}^{c} PDF_{Eqv_j}$$

where c represents the number of assemblages. This can easily extend to demonstrate the property of regularity, which states that if all the assemblages have the same diversity and are functionally dissimilar, then the global diversity can be calculated as the product of the diversity of the assemblage by the number of assemblages. In a $PDF_{Eqv}$ framework, if we assume that $\int \min\left\{max_{1\leq i\leq S_j}\{PDF_i\ (x)\ w_i\}, max_{1\leq i\leq S_k}\{PDF_i\ (x)\ w_i\}\right\}\, dx = 0$ for all $j \neq \mathrm{k}$, and $PDF_{Eqv_j} = PDF_{Eqv_k}$, then:

$$PDF_{Eq_\Upsilon} = \sum_{j=1}^{c} PDF_{Eqv_j} = c\ \times PDF_{Eqv_j}$$

**Method S1. Plant functional trait database**

We used data from 40 plots in a Mediterranean grassland located 20 km north of Madrid, in central Spain (40°360 N, 3°450 W; elevation ~ 700 m; Carmona et al. 2015). Plots were established along a topographic gradient extending from the upper slope, which is characterized by shallow soils and lower nutrient and water availability, to the lower slope, where soils are deeper and substantially more humid. Within each plot, 6 to 10 individuals were randomly selected for each of the most abundant species (i.e., those accounting for at least 90% of the total plot cover), resulting in a total of 2,540 individuals. For each individual two traits were measured: (1) plant height (cm), measured as the distance between the highest photosynthetic leaf and the plant's base; and (2) specific leaf area (SLA; mm2/mg), as the ratio between the surface of the leave and its dry weight (Pérez-Harguindeguy *et al.* 2013). These two traits, together, represent two different axes of aboveground trait variation in plants (Díaz *et al.* 2016).

### Method S2. Carnivora data curation and construction of climatic space

We downloaded 7,940,355 records from GBIF that has coordinates values, no geospatial issues assigned discarding fossils specimens. Then, we filtered out those records assigned as 'material sample', 'material citation' (mostly extinct species) and 99 'living specimen'. Within the taxonomic validation, we kept records identified at species, subspecies, form or variety taxon rank. We then check the status using Wilson & Reeder (2005) removing any doubtful species name and reassigning the accepted species name to those records that had synonyms (infraspecies were grouped into their corresponding accepted). Geographically, we removed capital and country centroids and records placed near biodiversity institutions (Zizka *et al.* 2019). We also validated records whose coordinates fall and match the country assigned and manually check records placed at borders (none was discarded). For those records that were placed in 'sea' areas, we assumed that they were valid if fell at <1km from coast for species that do not live in marine habitats (to overcome issues with shape and resolution of coasts in the shapefile used for the validation or rounded/trunk coordinates). For species of Phocidae, Otariidae y Odobenidae and *Enhydra lutris* & *Lontra felina* we kept records that were placed at < 50 km from the coast and we did not discard any record of *Ursus maritimus* as this species can be found on the ice and far away from the coast. The validated dataset (4,664,112) were simplified by removing duplicates as: (=) species + (=) coordinates + (=) date of collection + (=) institution]. The validated dataset includes 4,664,112 records, but those species with less than 30 records were excluded to guarantee a proper estimation of bioclimatic niches, resulting in a total of 250 species. The climatic niche of each species was described using six bioclimatic variables derived from CHELSA climate dataset version 2.1 (Brun *et al.* 2022) that were shown to be effective in modelling species' climatic preferences (Quintero & Wiens 2013): annual mean temperature (BIO1), maximum temperature of the warmest month (BIO5), minimum temperature of the coldest month (BIO6), annual precipitation (BIO12), precipitation of the wettest month (BIO13), and precipitation of the driest month (BIO14). We performed a principal component analysis on scaled versions of the bioclimatic variables to reduce the dimensionality of the data and run a Horn's parallel analysis, as implemented in the paran R package (Dinno 2009), and retained two components which together accounted for 83.78% of the variation (46.25 associated with the first component and 37.53% associated with the second component; see Figure 4a).

### Method S3. Atlantic cod data description and curation

Individuals were captured and tagged around mid-May 2012 using fyke-nets deployed at depths of 1–10 m and soaked for 1–3 days. Individuals were anesthetized with clove oil and surgically implanted with Innovasea V9P and V13P acoustic transmitters (delay = 110-250 sec, battery life = 350-1292 days) in the abdominal cavity following standard procedures (Villegas-Ríos *et al.* 2017). After full recovery from anaesthesia (generally within 5–10 min), fish were released at their capture sites (see Villegas-Ríos et al., 2016, 2021 for details). Cod movement within the fjord was monitored with 33 Innovasea VR2W omnidirectional receivers fixed at three-meter depth and pointing downwards, which provided a good coverage of the study area (Villegas-Ríos et al., 2016). To study changes along the year, we used data for 25 fish for which there were regular detections (i.e., no gaps longer than three days) between May 24$^{th}$ 2012 and June 5$^{th}$ 2013. Fish had an average size (measured as body length) of 49 cm, ranging from 31 to 75 cm. Detections were used to calculate centres of activity (COA) for each individual at regular 30 min time bins, as described in Villegas-Ríos *et al.* (2017).

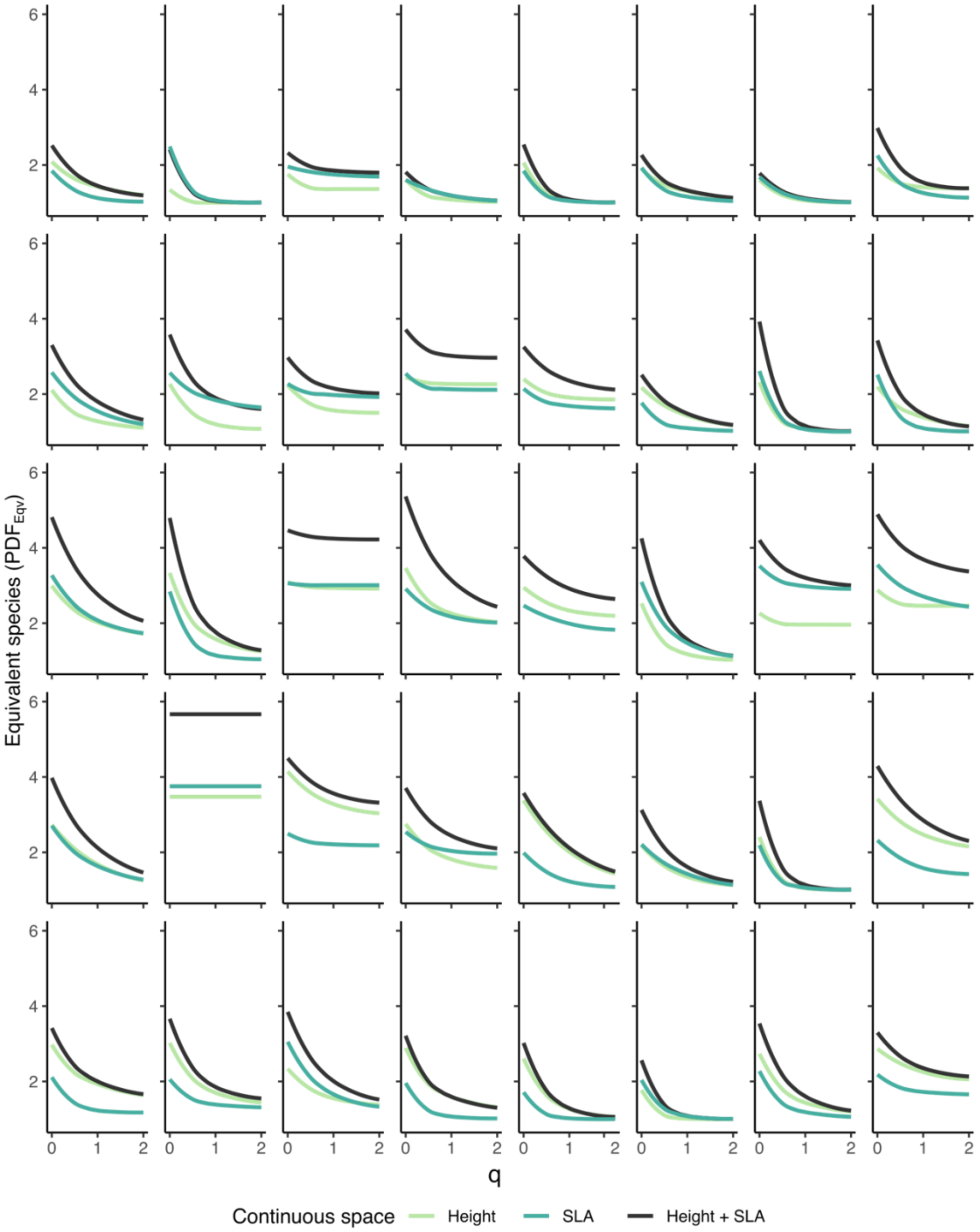


**Figure S1. Diversity profiles calculated for 40 plots along a topographic gradient in a Mediterranean grassland using three different continuous spaces.** The number of equivalent species was calculated as the equivalent probability density function number ($PDF_{Eqv}$) along values of the parameter q, which weights they importance of rare species in comparison with abundant species, ranging from 0 and 2. $PDF_{Eqv}$ was calculated using plant height (light green), specific leaf area (SLA; dark green) and the two-dimensional continuous space formed by height and SLA (black).

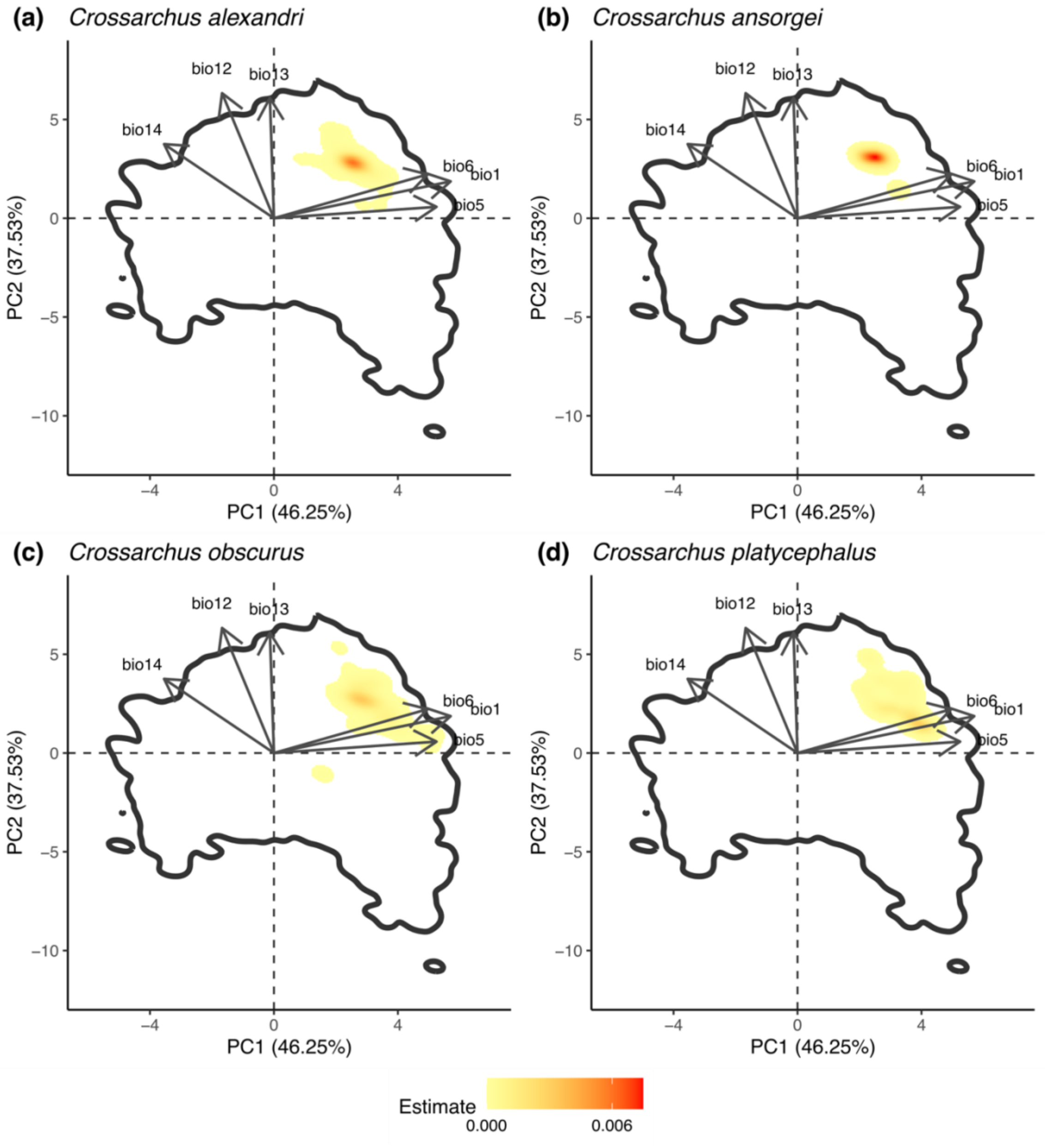


**Figure S2. Bioclimatic niche of species in the genus *Crossarchus*.** Probability density functions (PDFs) were used to calculate the bioclimatic niche by using two dimensions of a principal component analysis (PCA) based on six bioclimatic variables (see main text and Figure 4). Colours correspond to the estimate of the PDF function and black contour indicates the 0.99 quantile of the order Carnivora bioclimatic space calculated as the sum of the PDFs of all species.

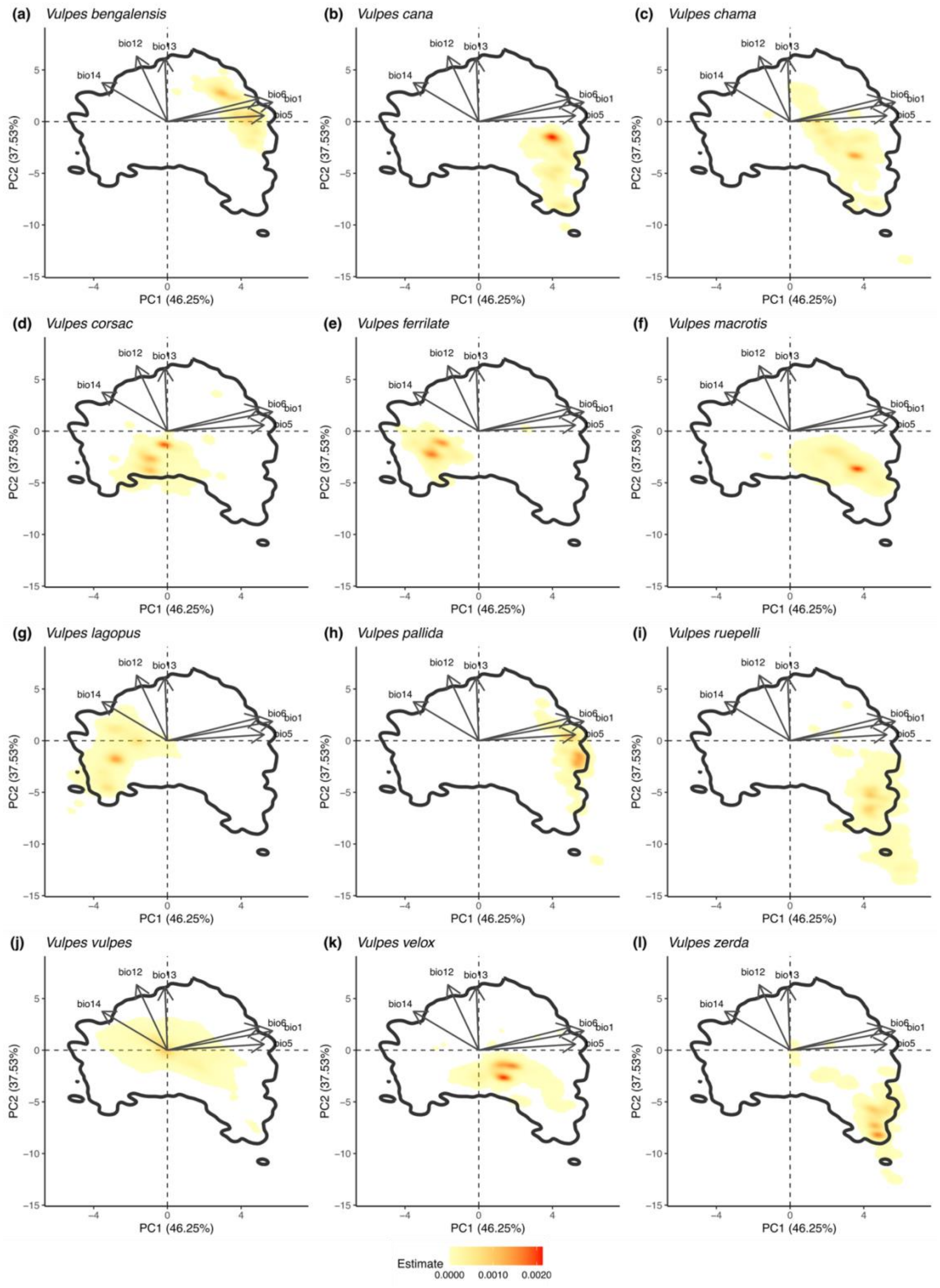


**Figure S3. Bioclimatic niche of species in the genus *Vulpes*.** Probability density functions (PDFs) were used to calculate the bioclimatic niche of each *Vulpes* species by using two dimensions of a principal component analysis (PCA) based on six bioclimatic variables (see main text and Figure 4). Colours correspond to the estimate of the PDF function and black contour indicates the 0.99 quantiles of the Carnivora bioclimatic space calculated as the sum of the PDFs of all species.

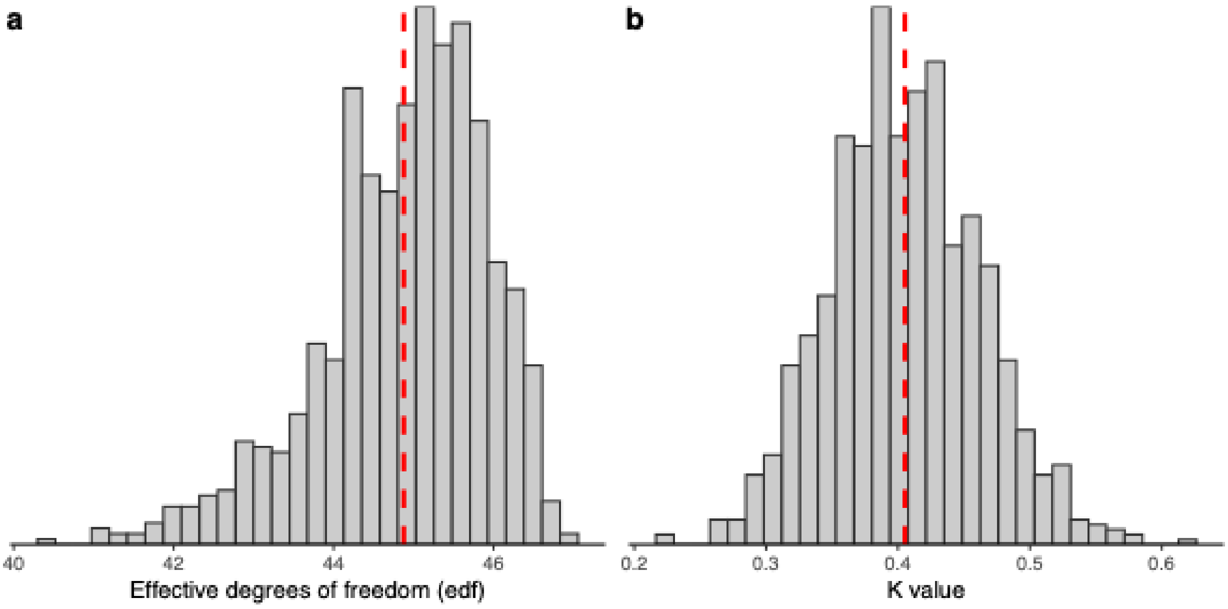


**Figure S4. Distribution of the (a) effective degrees of freedom and (b) K values for each of the bootstraps in the generalized additive model (GAM).** Red vertical lines indicate the mean value.

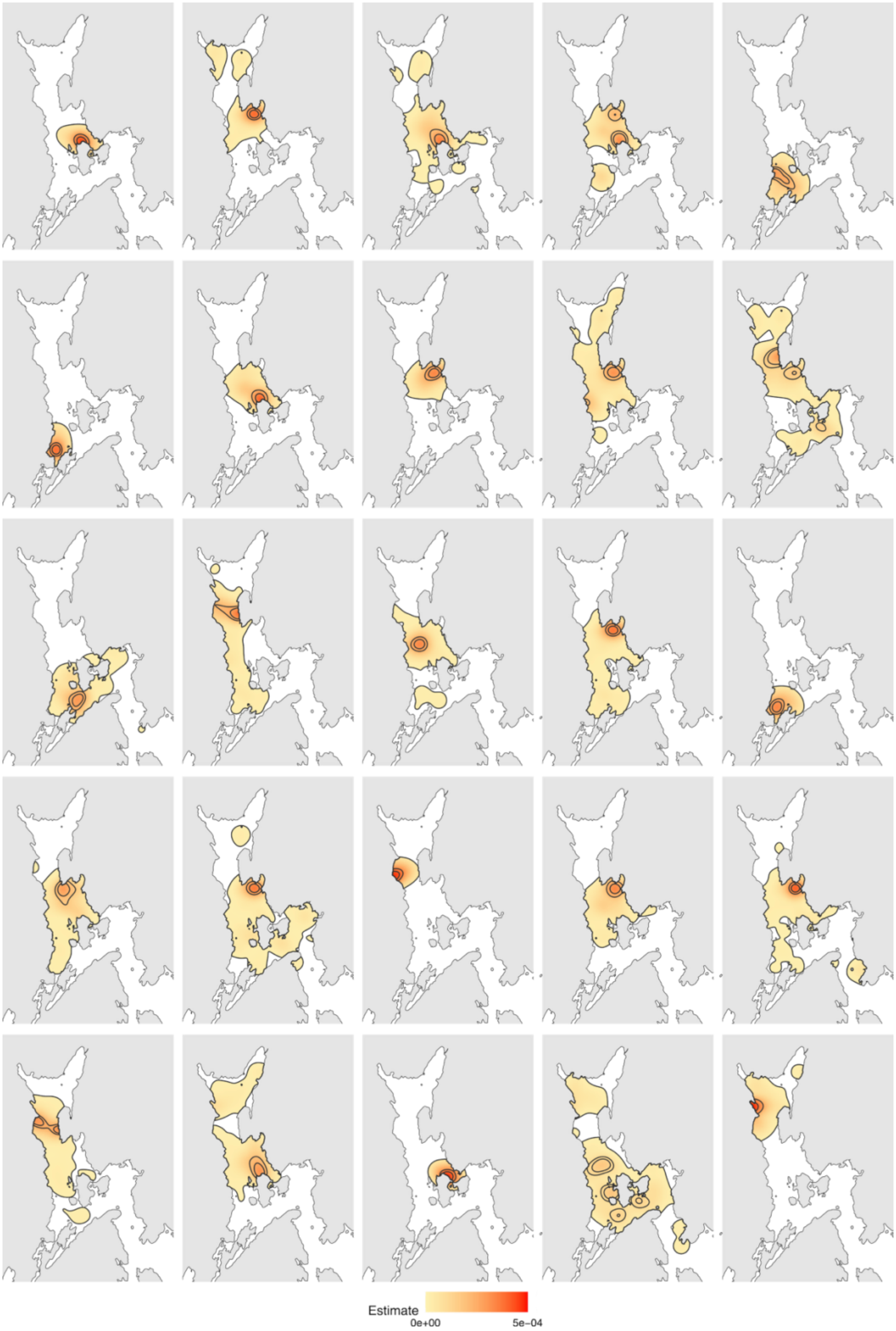


**Figure S5. Occupation of the geographic space by each individual in the period of one year.** Each panel represents the probability density function (PDF) estimated by using the centres of activity (COA) of a fish in the period of one year. Grey areas correspond to land while coloured areas represent the estimate of the PDF. Further, black isolines represent the 0.25, 0.5 and 0.99 quantiles.

**Table S1. Summary of post-hoc Tukey tests comparing the equivalent probability density function number ($PDF_{Eqv}$) calculated using three different trait spaces, two of them based on unique traits (height and specific leaf area (SLA)) and one being a multidimensional trait space formed by these two traits.** Comparisons were made for $PDF_{Eqv}$ calculated for different values of the parameter q, which weights the importance given to rare species compare to dominant species. A likelihood ratio test in linear mixed-effects models testing the effect of the trait space on $PDF_{Eqv}$ revealed strong significance ($p < 0.001$) for every value of q. Significant comparisons are highlighted in bold.

| q | χ2 (df = 2) | Comparison | Estimate | Standard error | z value | p-value (Tukey test) |
|---|---|---|---|---|---|---|
| 0 | 92.36 | SLA - Height | -0.11 | 0.09 | -1.232 | 0.43 |
| | | **Multitrait -Height** | **0.98** | **0.09** | **10.607** | **< 0.001** |
| | | **Multitrait - SLA** | **1.1** | **0.09** | **11.839** | **< 0.001** |
| 0.1 | 88.52 | SLA - Height | -0.12 | 0.09 | -1.34 | 0.37 |
| | | **Multitrait -Height** | **0.88** | **0.09** | **10.15** | **< 0.001** |
| | | **Multitrait - SLA** | **1** | **0.09** | **11.49** | **< 0.001** |
| 0.2 | 82.26 | SLA - Height | -0.12 | 0.08 | -1.4 | 0.34 |
| | | Multitrait -Height | 0.79 | 0.08 | **9.48** | **< 0.001** |
| | | Multitrait - SLA | 0.91 | 0.08 | **10.88** | **< 0.001** |
| 0.3 | 75.62 | SLA - Height | -0.11 | 0.08 | -1.42 | 0.33 |
| | | **Multitrait -Height** | **0.71** | **0.08** | **8.81** | **< 0.001** |
| | | **Multitrait - SLA** | **0.82** | **0.08** | **10.22** | **< 0.001** |
| 0.4 | 69.32 | SLA - Height | -0.11 | 0.08 | -1.4 | 0.34 |
| | | **Multitrait -Height** | **0.64** | **0.08** | **8.2** | **< 0.001** |
| | | **Multitrait - SLA** | **0.75** | **0.08** | **9.6** | **< 0.001** |
| 0.5 | 63.52 | SLA - Height | -0.1 | 0.08 | -1.38 | 0.35 |
| | | **Multitrait -Height** | **0.58** | **0.08** | **7.64** | **< 0.001** |
| | | **Multitrait - SLA** | **0.68** | **0.08** | **9.03** | **< 0.001** |
| 0.6 | 58.44 | SLA - Height | -0.1 | 0.08 | -1.37 | 0.36 |
| | | Multitrait -Height | 0.53 | 0.08 | **7.16** | **< 0.001** |
| | | Multitrait - SLA | 0.64 | 0.08 | **8.53** | **< 0.001** |
| 0.7 | 54.04 | SLA - Height | -0.1 | 0.07 | -1.35 | 0.36 |
| | | **Multitrait -Height** | **0.49** | **0.07** | **6.74** | **< 0.001** |
| | | **Multitrait - SLA** | **0.59** | **0.07** | **8.1** | **< 0.001** |
| 0.8 | 50.22 | SLA - Height | -0.09 | 0.07 | -1.33 | 0.38 |
| | | **Multitrait -Height** | **0.46** | **0.07** | **6.39** | **< 0.001** |
| | | **Multitrait - SLA** | **0.55** | **0.07** | **7.72** | **< 0.001** |
| 0.9 | 46.9 | SLA - Height | -0.09 | 0.07 | -1.3 | 0.39 |
| | | **Multitrait -Height** | **0.43** | **0.07** | **6.08** | **< 0.001** |
| | | **Multitrait - SLA** | **0.52** | **0.07** | **7.38** | **< 0.001** |
| 1 | 43.99 | SLA - Height | -0.09 | 0.07 | -1.26 | 0.42 |
| | | **Multitrait -Height** | **0.4** | **0.07** | **5.83** | **< 0.001** |
| | | **Multitrait - SLA** | **0.49** | **0.07** | **7.09** | **< 0.001** |
| 1.1 | 41.43 | SLA - Height | -0.08 | 0.07 | -1.21 | 0.44 |
| | | **Multitrait -Height** | **0.38** | **0.07** | **5.6** | **< 0.001** |

| | | | | | | |
|---|---|---|---|---|---|---|
| | | **Multitrait - SLA** | **0.46** | **0.07** | **6.82** | **< 0.001** |
| | | SLA - Height | -0.08 | 0.07 | -1.17 | 0.47 |
| 1.2 | 39.16 | **Multitrait -Height** | **0.36** | **0.07** | **5.4** | **< 0.001** |
| | | **Multitrait - SLA** | **0.44** | **0.07** | **6.579** | **< 0.001** |
| | | SLA - Height | -0.07 | 0.07 | -1.13 | 0.5 |
| 1.3 | 37.13 | **Multitrait -Height** | **0.34** | **0.07** | **5.23** | **< 0.001** |
| | | **Multitrait - SLA** | **0.42** | **0.07** | **6.36** | **< 0.001** |
| | | SLA - Height | -0.07 | 0.06 | -1.08 | 0.53 |
| 1.4 | 35.3 | **Multitrait -Height** | **0.33** | **0.06** | **5.08** | **< 0.001** |
| | | **Multitrait - SLA** | **0.4** | **0.06** | **6.16** | **< 0.001** |
| | | SLA - Height | -0.07 | 0.06 | -1.02 | 0.56 |
| 1.5 | 33.63 | **Multitrait -Height** | **0.32** | **0.06** | **4.94** | **< 0.001** |
| | | **Multitrait - SLA** | **0.38** | **0.06** | **5.97** | **< 0.001** |
| | | SLA - Height | -0.06 | 0.06 | -0.98 | 0.59 |
| 1.6 | 32.09 | **Multitrait -Height** | **0.31** | **0.06** | **4.82** | **< 0.001** |
| | | **Multitrait - SLA** | **0.37** | **0.06** | **5.79** | **< 0.001** |
| | | SLA - Height | -0.06 | 0.06 | -0.93 | 0.62 |
| 1.7 | 30.69 | **Multitrait -Height** | **0.3** | **0.06** | **4.7** | **< 0.001** |
| | | **Multitrait - SLA** | **0.36** | **0.06** | **5.63** | **< 0.001** |
| | | SLA - Height | -0.06 | 0.06 | -0.885 | 0.65 |
| 1.8 | 29.4 | **Multitrait -Height** | **0.29** | **0.06** | **4.6** | **< 0.001** |
| | | **Multitrait - SLA** | **0.35** | **0.06** | **5.48** | **< 0.001** |
| | | SLA - Height | -0.05 | 0.06 | -0.84 | 0.68 |
| 1.9 | 28.22 | **Multitrait -Height** | **0.28** | **0.06** | **4.5** | **< 0.001** |
| | | **Multitrait - SLA** | **0.33** | **0.06** | **5.34** | **< 0.001** |
| | | SLA - Height | -0.05 | 0.06 | -0.8 | 0.7 |
| 2 | 27.14 | **Multitrait -Height** | **0.28** | **0.06** | **4.41** | **< 0.001** |
| | | **Multitrait - SLA** | **0.33** | **0.06** | **5.21** | **< 0.001** |

**Table S2. Summary of post-hoc Tukey tests comparing the equivalent numbers (calculated using the Hill-Chao framework) with three different dissimilarity matrices, two of them based on unique traits (height and specific leaf area (SLA)) and one being the Gower's distance using these two traits.** Comparisons were made for equivalent numbers calculated for different values of the parameter q, which weights the importance given to rare species compare to dominant species.

| q | χ2 (df = 2) | p-value (likelihood ratio test) | Comparison | Estimate | Standard error | z value | p-value (Tukey test) |
|---|---|---|---|---|---|---|---|
| 0 | 0.44 | 0.8 | SLA - Height | -0.05 | 0.09 | -0.50 | 0.87 |
| | | | Multitrait -Height | -0.06 | 0.09 | -0.61 | 0.81 |
| | | | Multitrait - SLA | -0.01 | 0.09 | -0.11 | 0.99 |
| 0.1 | 0.5 | 0.78 | SLA - Height | -0.05 | 0.09 | -0.54 | 0.85 |
| | | | Multitrait -Height | -0.06 | 0.09 | -0.66 | 0.79 |
| | | | Multitrait - SLA | -0.01 | 0.09 | -0.11 | 0.99 |
| 0.2 | 0.56 | 0.75 | SLA - Height | -0.05 | 0.08 | -0.58 | 0.83 |
| | | | Multitrait -Height | -0.06 | 0.08 | -0.69 | 0.77 |
| | | | Multitrait - SLA | -0.01 | 0.08 | -0.11 | 0.99 |
| 0.3 | 0.61 | 0.74 | SLA - Height | -0.05 | 0.08 | -0.61 | 0.82 |
| | | | Multitrait -Height | -0.06 | 0.08 | -0.72 | 0.75 |
| | | | Multitrait - SLA | -0.01 | 0.08 | -0.11 | 0.99 |
| 0.4 | 0.64 | 0.72 | SLA - Height | -0.05 | 0.08 | -0.63 | 0.81 |
| | | | Multitrait -Height | -0.06 | 0.08 | -0.73 | 0.74 |
| | | | Multitrait - SLA | -0.01 | 0.08 | -0.11 | 0.99 |
| 0.5 | 0.67 | 0.72 | SLA - Height | -0.05 | 0.07 | -0.64 | 0.80 |
| | | | Multitrait -Height | -0.06 | 0.07 | -0.75 | 0.73 |
| | | | Multitrait - SLA | -0.01 | 0.07 | -0.11 | 0.99 |
| 0.6 | 0.68 | 0.71 | SLA - Height | -0.05 | 0.07 | -0.64 | 0.80 |
| | | | Multitrait -Height | -0.06 | 0.07 | -0.76 | 0.73 |
| | | | Multitrait - SLA | -0.01 | 0.07 | -0.11 | 0.99 |
| 0.7 | 0.69 | 0.71 | SLA - Height | -0.05 | 0.07 | -0.65 | 0.79 |
| | | | Multitrait -Height | -0.05 | 0.07 | -0.76 | 0.73 |
| | | | Multitrait - SLA | -0.01 | 0.07 | -0.12 | 0.99 |
| 0.8 | 0.69 | 0.71 | SLA - Height | -0.05 | 0.07 | -0.64 | 0.80 |
| | | | Multitrait -Height | -0.05 | 0.07 | -0.76 | 0.73 |
| | | | Multitrait - SLA | -0.01 | 0.07 | -0.12 | 0.99 |
| 0.9 | 0.68 | 0.71 | SLA - Height | -0.04 | 0.07 | -0.64 | 0.80 |
| | | | Multitrait -Height | -0.05 | 0.07 | -0.76 | 0.73 |
| | | | Multitrait - SLA | -0.01 | 0.07 | -0.13 | 0.99 |
| 1 | 0.67 | 0.71 | SLA - Height | -0.04 | 0.07 | -0.64 | 0.80 |
| | | | Multitrait -Height | -0.05 | 0.07 | -0.76 | 0.73 |
| | | | Multitrait - SLA | -0.01 | 0.07 | -0.13 | 0.99 |
| 1.1 | 0.66 | 0.71 | SLA - Height | -0.04 | 0.07 | -0.62 | 0.81 |
| | | | Multitrait -Height | -0.05 | 0.07 | -0.76 | 0.73 |
| | | | Multitrait - SLA | -0.01 | 0.07 | -0.14 | 0.99 |

| | | | | | | | |
|---|---|---|---|---|---|---|---|
| | | | SLA - Height | -0.04 | 0.07 | -0.61 | 0.82 |
| 1.2 | 0.65 | 0.72 | Multitrait -Height | -0.05 | 0.07 | -0.75 | 0.73 |
| | | | Multitrait - SLA | -0.01 | 0.07 | -0.15 | 0.99 |
| | | | SLA - Height | -0.04 | 0.06 | -0.59 | 0.82 |
| 1.3 | 0.64 | 0.72 | Multitrait -Height | -0.05 | 0.06 | -0.75 | 0.73 |
| | | | Multitrait - SLA | -0.01 | 0.06 | -0.16 | 0.99 |
| | | | SLA - Height | -0.04 | 0.06 | -0.58 | 0.83 |
| 1.4 | 0.63 | 0.73 | Multitrait -Height | -0.05 | 0.06 | -0.74 | 0.74 |
| | | | Multitrait - SLA | -0.01 | 0.06 | -0.16 | 0.99 |
| | | | SLA - Height | -0.04 | 0.06 | -0.57 | 0.84 |
| 1.5 | 0.61 | 0.73 | Multitrait -Height | -0.05 | 0.06 | -0.74 | 0.74 |
| | | | Multitrait - SLA | -0.01 | 0.06 | -0.17 | 0.98 |
| | | | SLA - Height | -0.03 | 0.06 | -0.55 | 0.84 |
| 1.6 | 0.6 | 0.74 | Multitrait -Height | -0.05 | 0.06 | -0.74 | 0.74 |
| | | | Multitrait - SLA | -0.01 | 0.06 | -0.18 | 0.98 |
| | | | SLA - Height | -0.03 | 0.06 | -0.54 | 0.85 |
| 1.7 | 0.59 | 0.74 | Multitrait -Height | -0.05 | 0.06 | -0.73 | 0.74 |
| | | | Multitrait - SLA | -0.01 | 0.06 | -0.19 | 0.98 |
| | | | SLA - Height | -0.03 | 0.06 | -0.53 | 0.86 |
| 1.8 | 0.58 | 0.75 | Multitrait -Height | -0.04 | 0.06 | -0.73 | 0.75 |
| | | | Multitrait - SLA | -0.01 | 0.06 | -0.20 | 0.98 |
| | | | SLA - Height | -0.03 | 0.06 | -0.51 | 0.86 |
| 1.9 | 0.57 | 0.75 | Multitrait -Height | -0.04 | 0.06 | -0.73 | 0.75 |
| | | | Multitrait - SLA | -0.01 | 0.06 | -0.21 | 0.98 |
| | | | SLA - Height | -0.03 | 0.06 | -0.50 | 0.87 |
| 2 | 0.56 | 0.75 | Multitrait -Height | -0.04 | 0.06 | -0.72 | 0.75 |
| | | | Multitrait - SLA | -0.01 | 0.06 | -0.22 | 0.97 |
